# Transition from environmental to partial genetic sex determination in *Daphnia* through the evolution of a female-determining incipient W-chromosome

**DOI:** 10.1101/064311

**Authors:** Céline M.O. Reisser, Dominique Fasel, Evelin Hürlimann, Marinela Dukič, Cathy Haag-Liautard, Virginie Thuillier, Yan Galimov, Christoph Haag

## Abstract

Sex chromosomes can evolve during the evolution of genetic sex determination (GSD) from environmental sex determination (ESD). Despite theoretical attention, early mechanisms involved in the transition from ESD to GSD have yet to be studied in nature. No mixed ESD-GSD animal species have been reported, except for some species of *Daphnia*, small freshwater crustaceans in which sex is usually determined solely by the environment, but in which a dominant female sex-determining locus is present in some populations. This locus follows Mendelian single-locus inheritance, but has otherwise not been characterized genetically. We now show that the sex-determining genomic region maps to the same low-recombining peri-centromeric region of linkage group 3 (LG3) in three highly divergent populations of *D. magna*, and spans 3.6 Mb. Despite low levels of recombination, the associated region contains signs of historical recombination, suggesting a role for selection acting on several genes thereby maintaining linkage disequilibrium among the 36 associated SNPs. The region carries numerous genes involved in sex differentiation in other taxa, including *transformer2* and *sox9*. Taken together, the region determining the NMP phenotype shows characteristics of a sex-related supergene, suggesting that LG3 is potentially an incipient W chromosome despite the lack of significant additional restriction of recombination between Z and W. The occurrence of the female-determining locus in a pre-existing low recombining region illustrates one possible form of recombination suppression in sex chromosomes. *D. magna* is a promising model for studying the evolutionary transitions from ESD to GSD and early sex chromosome evolution.

## Introduction

Sex chromosomes have evolved independently multiple times in many taxa (Miura et al. 2008; Pokorná and Kratochvíl 2009; Stöck et al. 2011; Cheng et al. 2013; Tree of Sex Consortium 2014). Two evolutionary routes are thought to lead to the evolution of genetic sex determination, and sometimes of sex chromosomes. First, separate sexes may evolve from hermaphroditism, most likely through a male-sterility (i.e. female determining) mutation creating a breeding system called gynodioecy, with genetic females co-occurring with functional hermaphrodites. An initial male-determining mutation is also possible, but is less likely (Charlesworth and Charlesworth 1978). The sex-determining mutation can be favoured through causing obligate outcrossing, if inbreeding depression is severe, or if there is a fitness disadvantage to investing resources in both male and female functions compared to investing in only one sex (Charlesworth and Charlesworth 1978; Innes and Dunbrack 1993). Additional mutations in genes linked to the sex-determining locus may then be favoured if their effects are sexually antagonistic (i.e. have opposite effects in the two sexes; Rice 1987; Ellegren 2011). In the case of a female sex-determining locus, additional mutations include ones benefiting males, and deleterious in females (including female-sterility mutations creating males), which will often be eliminated unless they are linked to the femaleness-determining gene. Closer linkage will then be favored in this genomic region (Bull, 2006), which may lead to suppression of recombination. Eventually, this may result in a system with “proto-sex chromosomes” carrying linked genes that determine both sexes genetically (with male and female determining mutations on opposite homologs).

The second route towards evolving GSD and sex chromosomes may start from environmental sex determination (ESD), which could be the ancestral state in several major animal groups (Ohno 1967; Pokorná and Kratochvíl 2016). Although transition from ESD to GSD may be gradual, involving shifting genotype-specific thresholds for male vs. female development under fluctuating environmental conditions (Van Dooren and Leimar 2003), a scenario similar to that of the transition from hermaphroditism to separate sexes is also plausible: a female-determining mutation in a population with pure ESD could lead to a system in which females are genetically determined, while other individuals have ESD (the route through an initial male-determining mutation is also possible). Such a sex-determining mutation can be favoured if it restores a 1:1 sex ratio of the population, for example, after environmental conditions change (Edwards 1998), or through other advantages like those outlined for the first pathway above.

The evolution of suppressed recombination between sex chromosomes and more generally, the early stages of sex chromosome evolution remain poorly understood (Wright et al. 2016). Because of their intermediate position between hermaphroditic and dioecious species, gynodioecious plants might have been good candidates to study these early stages. However, they proved not to be very informative in this respect, because gynodioecy is often controlled by cyto-nuclear interactions between mitochondrial male sterility mutations and nuclear “restorer” genes (McCauley and Bailey 2009; Beukeboom and Perrin 2014). Only a few species appear to have pure nuclear control and may therefore conform to the above theory of sex chromosome evolution (Kohn 1988; Connor and Charlesworth 1989; Weller and Sakai 1991; Spigler et al. 2008). Species in transition from ESD to GSD are therefore of particular interest, although the transition may be rapid (Pokorná and Kratochvíl 2009), so that few systems are available for study.

In some populations of *Daphnia*, pure ESD individuals co-occur with genetically determined females (Innes and Dunbrack 1993, Tessier and Caceres 2004, Galimov et al. 2011). *Daphnia* are cyclical parthenogens, in which clonal reproduction with live born offspring is interspersed with sexual reproduction phases producing diapause stages. Sex determination is usually environmental (Hobaek and Larson 1990, Zhang and Baer 2000, LeBlanc and Medlock 2015), with cues differing among populations (Roulin et al. 2013). In nature, male development of the clonal offspring present in the ovaries may be elicited by the mother emitting a juvenile hormone (JH) or a JH pathway-related molecule (Olmstead and Leblanc 2002). Male production can also be artificially induced by adding hormone analogs to the culture medium (Olmstead and Leblanc 2002). However, some *Daphnia* individuals never produce males, neither under natural conditions nor when artificially exposed to hormone analogs (Innes and Dunbrack 1993; Tessier and Caceres 2004; Galimov et al. 2011). This non-male-producer “NMP” trait segregates as a single Mendelian locus (or single region) with a dominant female-determining allele called “W”. Heterozygous genotypes (“ZW”) are genetically determined females called NMP individuals, which take part in sexual diapause stage production exclusively as females. Homozygous genotypes (“ZZ”) are cyclical parthenogens with ESD. They produce diapause stages either through male or female function, and are called male producers (“MP” individuals; Galimov et al. 2011).

*Daphnia* populations harboring NMP as well as MP individuals offer an opportunity to study the transition from ESD to GSD, and potentially the early stages of sex chromosome evolution. At present, female is the only genetically determined sex in the populations, so that the situation resembles gynodioecy, with no genetically determined males present. Nonetheless, the early steps of sex chromosome evolution may have taken place on both homologs. For instance, sexually antagonistic mutations may have occurred in genes linked to the female-determining locus, and suppressed recombination may have evolved in response to such two-locus polymorphisms in the region.

We used genetic linkage mapping, association mapping, and comparative genomics methods to genetically characterize the sex-determining locus and its surrounding region in *D. magna*. Specifically, we first investigated whether the sex-determining locus maps to the same genomic location in crosses involving NMP females from populations with highly divergent mitochondrial lineages (Galimov et al. 2011, Svendsen et al. 2015). This is relevant because the W allele is expected to get co-transmitted with the mitochondrial haplotype (as both have female-limited transmission). Hence the existence of NMP in these divergent lineages might be due to recurrent evolution of a W allele, or, alternatively, might hint at long-term maintenance of the polymorphism within this species. The recombinant frequencies among the offspring of the experimental crosses were also used to investigate the hypothesis of reduced recombination between W and Z. Second, we used association mapping within a single population to identify female-associated SNPs and test if the association among SNPs is maintained purely through physical linkage or also by selection. For this, we estimated levels of linkage disequilibrium and searched for traces of historical recombination between different associated SNPs in the region around the sex-determining locus. Finally, we screened the NMP associated region of *D. magna* for genes involved in sex-determination in this or other species to identify potential candidate female determining genes. The overall aim of these approaches was to gain a genetic understanding of the transition from ESD to GSD and to assess the possible evolution of an incipient ZW sex chromosome system in *D. magna*.

## Results

### Genetic linkage mapping of the NMP-determining region using microsatellites markers

We performed three NMPxMP crosses involving NMP females from three populations with highly divergent mitochondrial haplotypes. In each cross, F1 lines were phenotyped (MP vs. NMP) and genotyped at a total of 81 microsatellite loci to perform linkage mapping of the genomic regions underlying the NMP phenotype. The resulting maps were compared to the *Daphnia* genetic map (Dukič et al. in press), which is based on an F2 panel of an MPxMP cross.

In the NMPxMP_1 and NMPxMP_2 crosses, partially overlapping sets of thirteen markers were significantly linked to the NMP phenotype (or to other markers linked to it). In each cross, eleven of these markers were successfully genotyped in >70% of offspring and were thus used for mapping (see Supplementary Material S1). They all mapped to linkage group 3 (LG3) of the reference genetic map (Fig. 1). In the NMPxMP_1 cross, three markers (dm_scf02569_310402, dm_scf00933_2550 and dm_scf00700_81490) were completely linked with the NMP phenotype, as were five markers in the NMPxMP_2 cross (dm_scf02569_317703, dm_scf01492_1407, dm_scf00933_2550, dm_scf03156_57375 and dm_scf00966_75426). Only two of these markers were genotyped in the NMPxMP_3 cross, but they too showed complete linkage with the NMP phenotype (a third tested marker was not polymorphic in this cross, see Supplementary Material S1). In all three crosses, the fully linked markers mapped between cM positions 87.8 cM and 94.0 cM of LG3 in the reference genetic map. This region also contains the centromere (at 90.8 cM). We call this region the “NMP region” (Fig. 2). A Marey map of LG3 (Dukič et al. in press) shows that the NMP-region corresponds to a large (∼3 Mb) non-recombining region around the centromere (Fig. 2). Non-recombining regions around the centromeres are found on all linkage groups of *D. magna*, and are not a sign of reduced recombination between sex chromosomes, since they also occur on all other linkage groups in the MPxMP cross of the reference genetic map.

**Figure 1.**
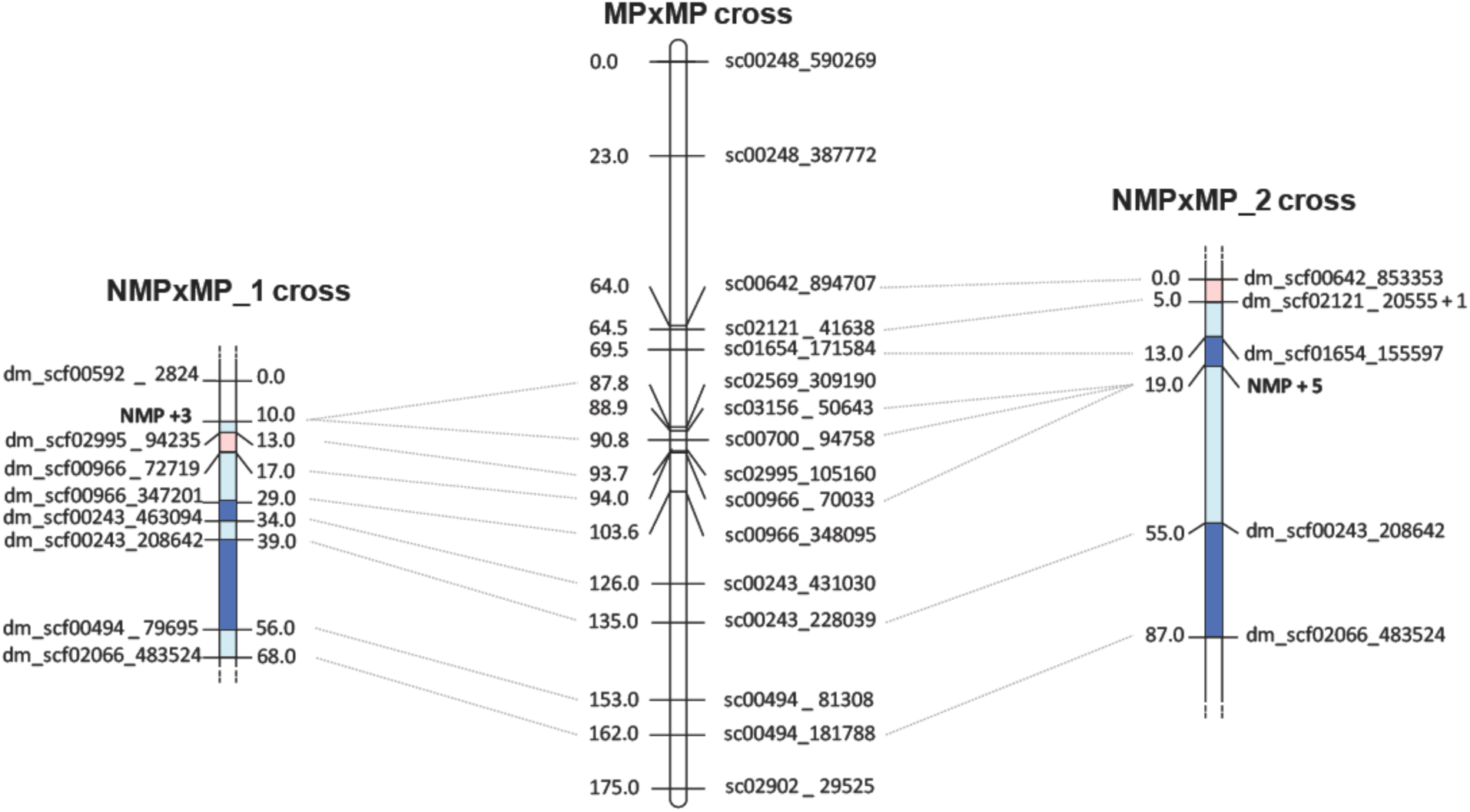
Genetic map of the two NMPxMP crosses (microsatellite markers) and of the MPxMP cross used to create the *Daphnia magna* reference genetic map (SNP markers, Dukič et al. in revision). Only Linkage Group 3 (LG3) is shown. Map distances are in centiMorgans, calculated with the Kosambi mapping function in R/qtl. Areas in light blue / light red show a non-significant reduction / expansion of recombination by comparison to the MP cross. while areas in bright blue indicates a significant reduction of recombination. For the two NMPxMP crosses, one marker per position is represented. In NMPxMP_1, NMP+3 indicates that three markers were in full linkage with the NMP locus. In NMPxMP_2, dm_scf00532_1398 was fully linked with dm_scf02121_20555; also, NMP+5 indicates that five markers were fully linked with the NMP locus.

**Figure 2.**
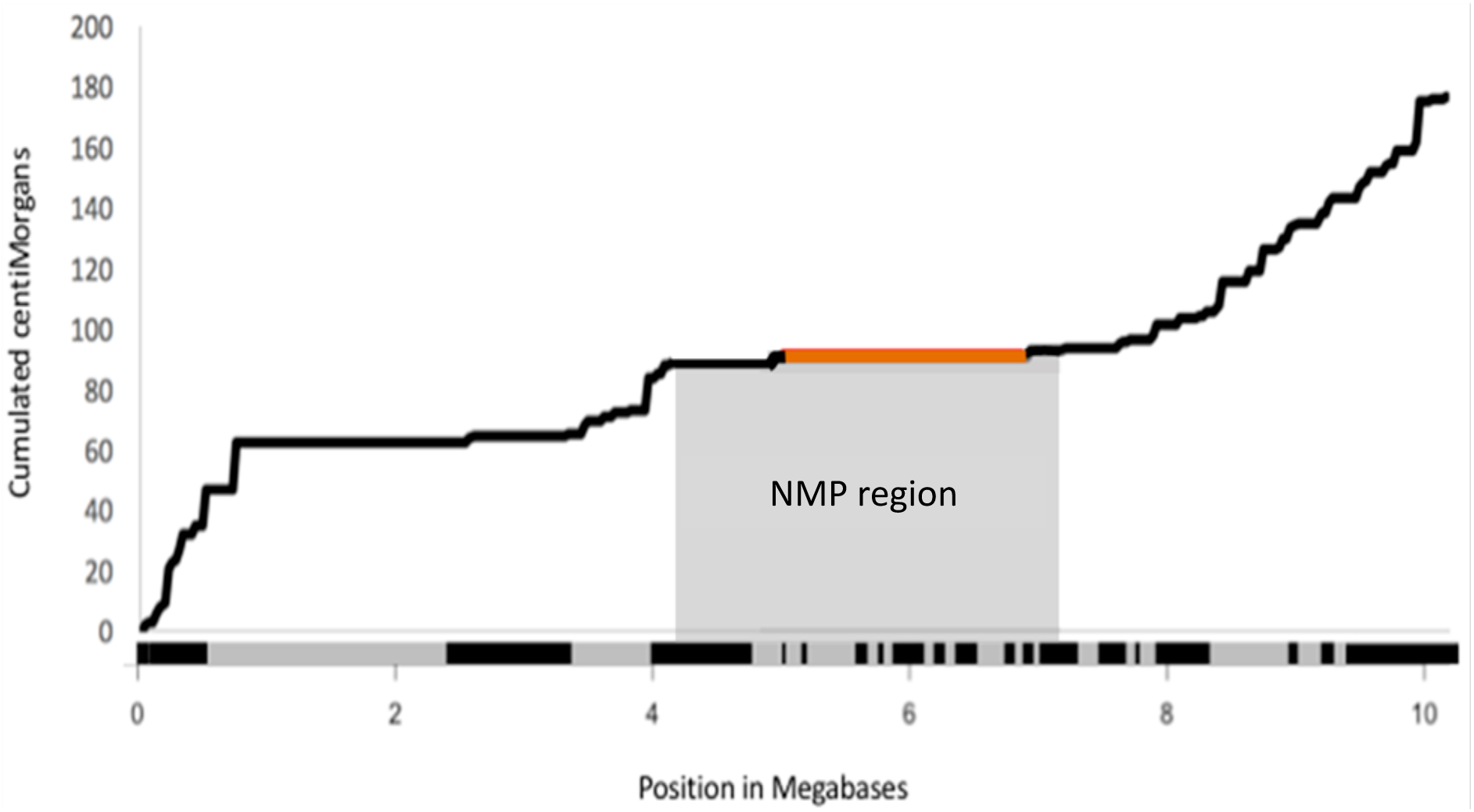
Marey map of LG3 in the MPxMP cross, showing the evolution of the recombination rate along the physical position on LG3. The x axis shows the position in Megabases. Alternated grey and black colors on the x axis represent the length of the different scaffolds that compose LG3. The centromeric region (90.8cM) is highlighted in orange. The shaded area corresponds to the genomic region containing fully linked markers with the NMP phenotype.

We investigated the possibility of additional recombination suppression in NMPxMP crosses as an indication of reduced recombination between the Z and (putative) W chromosomes compared to between two Z chromosomes. Indeed, genetic map distances between adjacent markers in and around the NMP region were on average lower in the NMPxMP crosses than in the reference genetic map (Fig. 1). However, this pattern is not statistically supported. The reduction was significant only for one, non-identical interval in each of the two crosses. Furthermore, the two crosses are not entirely independent. Both involved, at a different stage of the experiment, outcrossing of an NMP-female (different in the two crosses) to a male from the same distant population. In one cross this was done to create the actual mapping population, in the other it was done one generation earlier to obtain a highly heterozygous, hybrid NMP female with the intention to increase the number of markers available for mapping the W. Moreover, it is possible that the slight reduction in recombination frequencies was not specific to the NMP region (we did not have sufficient genotype data on linked markers in other genomic regions to test for this possibility). Overall, our results are clearly inconsistent with strongly reduced recombination across large parts of the putative incipient Z and incipient W chromosomes (Table 1), suggesting that if such a reduction has happened (compared to recombination in MPxMP crosses), it concerns only a small portion of the chromosome.

**Table 1.**
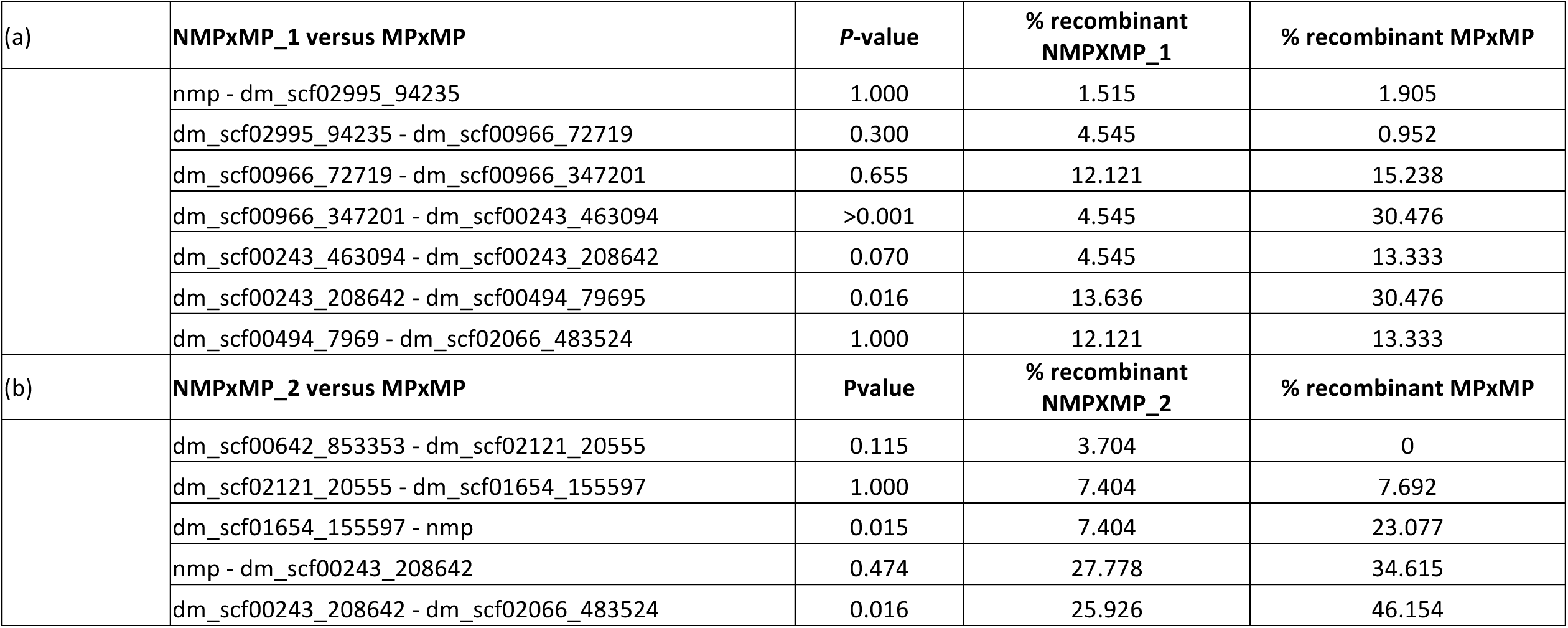
Comparison between crosses with one parent heterozygous (NMP x MP) and both parents homozygous for the MP factor. The table shows Fisher’s Exact Test *P*-values for all pairs of markers studied in (a) the NMPxMP_1 cross and (b) the NMPxMP_2 cross. Table also gives the proportion of recombinants for each cross, and for the reference cross (MPxMP, Dukič et al. in revision).

### Association mapping of SNPs in the NMP region

A genome-wide association analysis using RAD-sequencing of 72 individuals from a single population revealed 43 SNPs significantly associated (FDR <10^−5^) with the NMP phenotype (Fig. 3a, Supplementary material S2). Of these, 36 were contained in a region of LG3 between 72.3 and 95.7 cM, which is only slightly larger than the NMP region defined above (Fig. 3b). Of the other seven significantly associate SNPs, five mapped to two scaffolds on LG1, and one SNP to each of LG2 and LG4 (Table 2; Fig. 3). The number of associated SNPs per scaffold did not correlate with the scaffold size (Pearson’s correlation coefficient *r*^2^ = − 0.117). On the physical map of LG3 (see Methods and Dukič et al. in press), the main associations were distributed across 3.6 Mb, between positions 5.4 and 9.0 (Fig. 4.c; LG3 is 10.1Mb in total). The 36 highly associated SNPs are distributed across 15 non-contiguous scaffolds whose combined length is 2.42 Mb (about a quarter of LG3’s size). Thus, even though the physical map based on LD mapping may be incorrect, there is strong evidence that significantly associated SNPs are distributed across a large proportion of LG3. Interestingly, 25% of the significantly associated SNPs occurred on just a single 400kb scaffold. This scaffold, scf02569, is therefore a good candidate for the location of the initial, female-determining mutation.

**Table 2.**
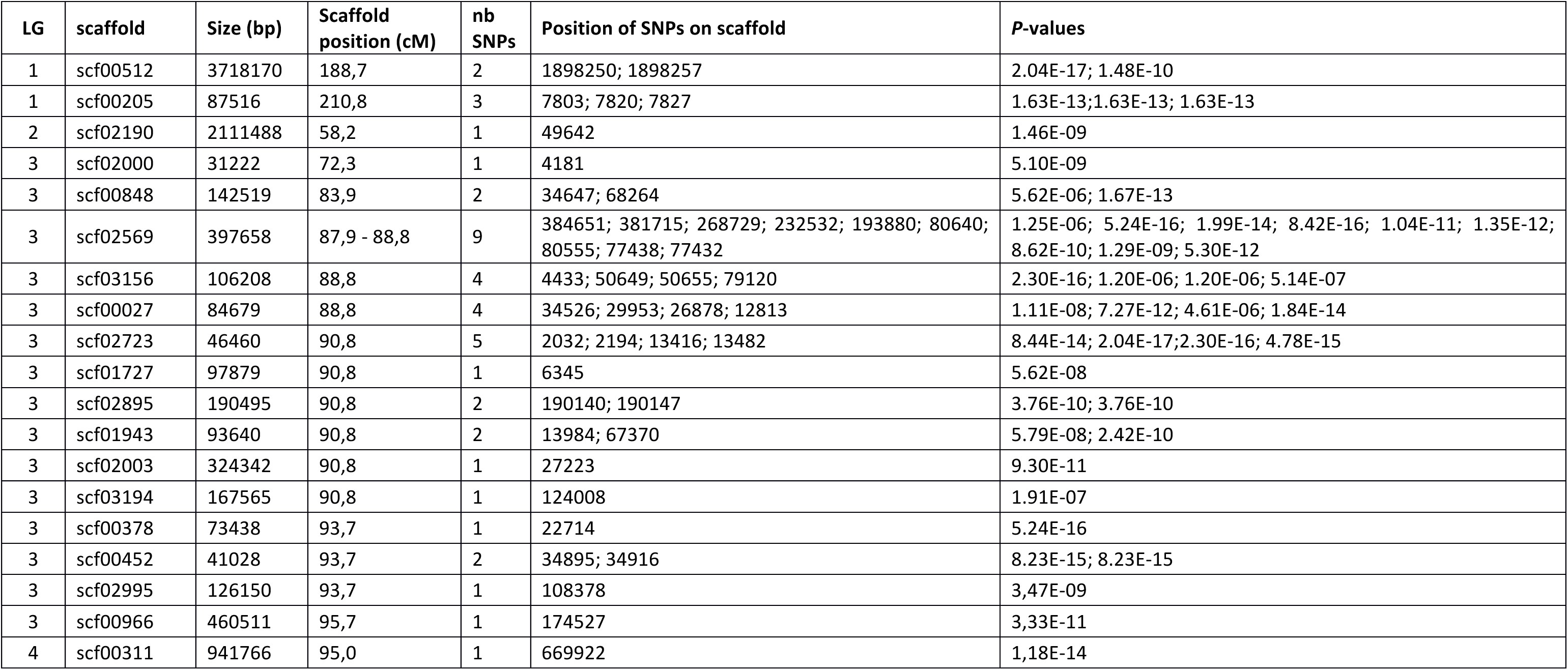
List of scaffolds containing SNPs significantly associated to NMP (Chi square test; *P* < 10^−5^). LG: linkage group; Size: total size of the scaffold (in basepair); nb. SNPs: number of associated SNPs on the scaffold; *P*-values: resulting Chi square P value.

| LG | scaffold | Size (bp) | Scaffold position (cM) | nb SNPs | Position of SNPs on scaffold | $P$-values |
| --- | --- | --- | --- | --- | --- | --- |
| 1 | scf00512 | 3718170 | 188,7 | 2 | 1898250; 1898257 | 2.04E-17; 1.48E-10 |
| 1 | scf00205 | 87516 | 210,8 | 3 | 7803; 7820; 7827 | 1.63E-13; 1.63E-13; 1.63E-13 |
| 2 | scf02190 | 2111488 | 58,2 | 1 | 49642 | 1.46E-09 |
| 3 | scf02000 | 31222 | 72,3 | 1 | 4181 | 5.10E-09 |
| 3 | scf00848 | 142519 | 83,9 | 2 | 34647; 68264 | 5.62E-06; 1.67E-13 |
| 3 | scf02569 | 397658 | 87,9 - 88,8 | 9 | 384651; 381715; 268729; 232532; 193880; 80640; 80555; 77438; 77432 | 1.25E-06; 5.24E-16; 1.99E-14; 8.42E-16; 1.04E-11; 1.35E-12; 8.62E-10; 1.29E-09; 5.30E-12 |
| 3 | scf03156 | 106208 | 88,8 | 4 | 4433; 50649; 50655; 79120 | 2.30E-16; 1.20E-06; 1.20E-06; 5.14E-07 |
| 3 | scf00027 | 84679 | 88,8 | 4 | 34526; 29953; 26878; 12813 | 1.11E-08; 7.27E-12; 4.61E-06; 1.84E-14 |
| 3 | scf02723 | 46460 | 90,8 | 5 | 2032; 2194; 13416; 13482 | 8.44E-14; 2.04E-17; 2.30E-16; 4.78E-15 |
| 3 | scf01727 | 97879 | 90,8 | 1 | 6345 | 5.62E-08 |
| 3 | scf02895 | 190495 | 90,8 | 2 | 190140; 190147 | 3.76E-10; 3.76E-10 |
| 3 | scf01943 | 93640 | 90,8 | 2 | 13984; 67370 | 5.79E-08; 2.42E-10 |
| 3 | scf02003 | 324342 | 90,8 | 1 | 27223 | 9.30E-11 |
| 3 | scf03194 | 167565 | 90,8 | 1 | 124008 | 1.91E-07 |
| 3 | scf00378 | 73438 | 93,7 | 1 | 22714 | 5.24E-16 |
| 3 | scf00452 | 41028 | 93,7 | 2 | 34895; 34916 | 8.23E-15; 8.23E-15 |
| 3 | scf02995 | 126150 | 93,7 | 1 | 108378 | 3,47E-09 |
| 3 | scf00966 | 460511 | 95,7 | 1 | 174527 | 3,33E-11 |
| 4 | scf00311 | 941766 | 95,0 | 1 | 669922 | 1,18E-14 |

**Figure 3.**
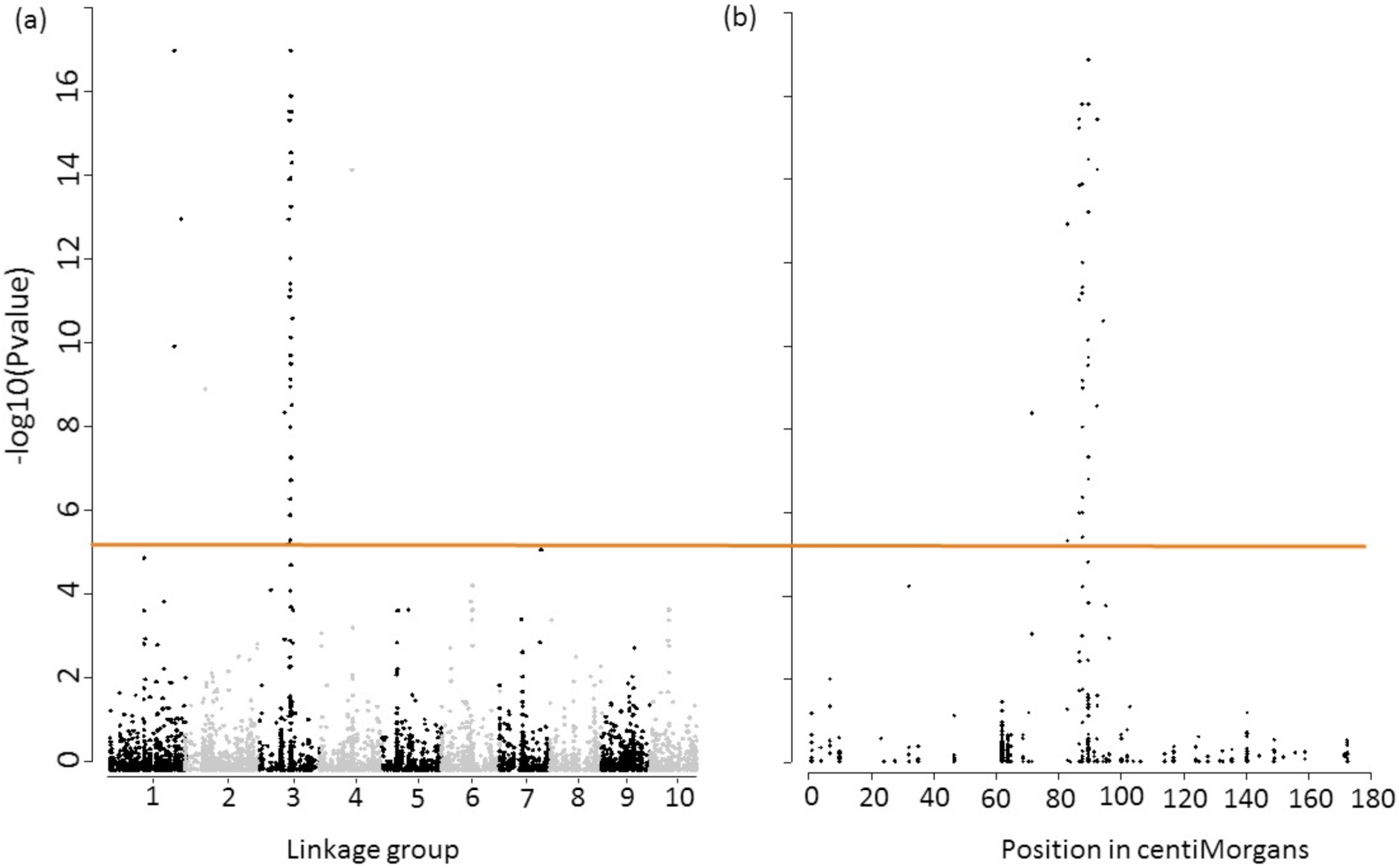
Genome-wide association results. Associations of SNP loci with the NMP polymorphism in a sample of 53 MP and 17 NMP individuals (a) across the entire genome and (b) on LG3. On LG3, markers between 72.3cM and 95.7cM show significant associations with the NMP phenotype (the orange line shows significance, with FDR-corrected P-values < 10^−3^).

**Figure 4.**
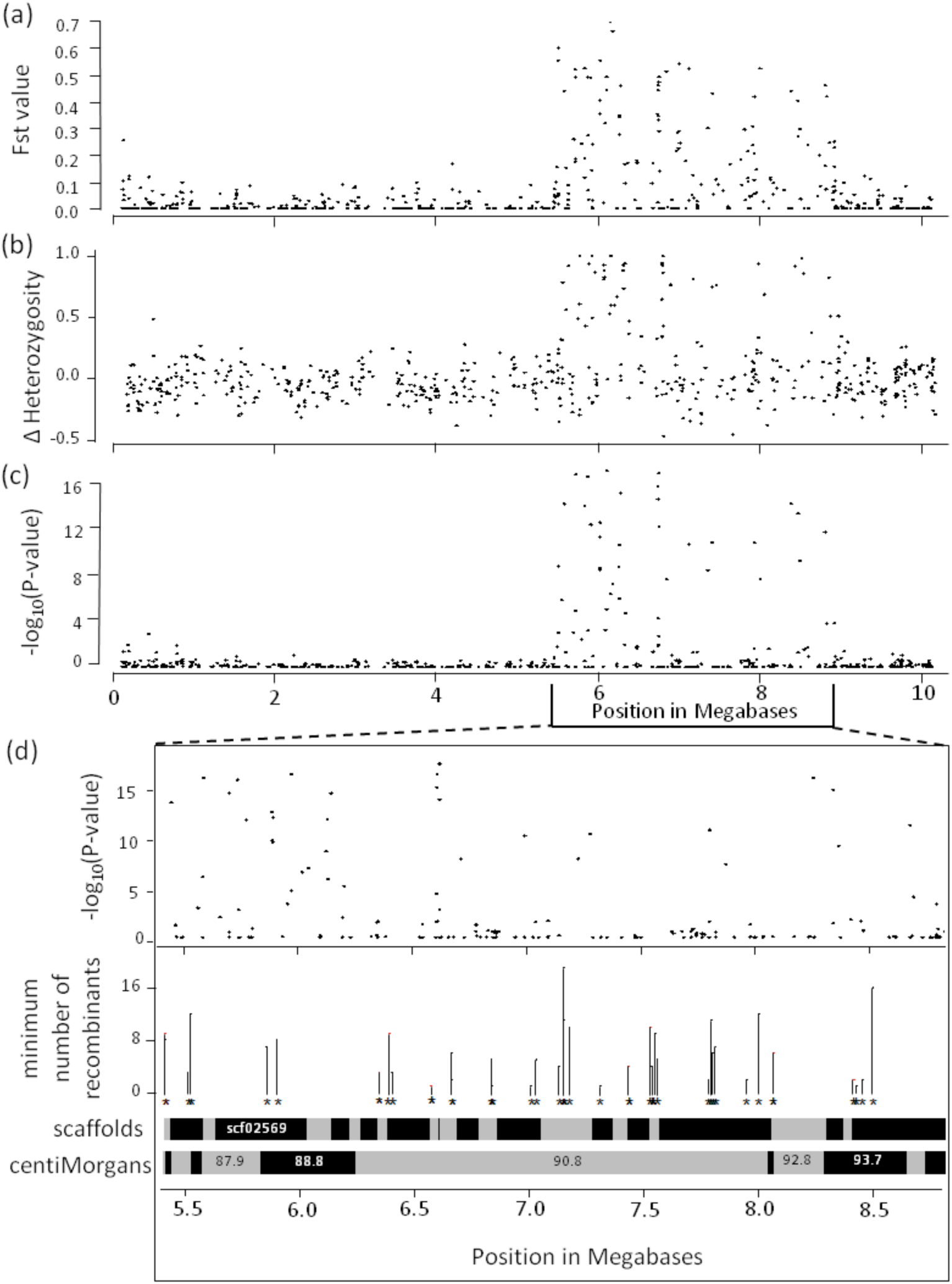
Subdivision between NMP and MP individuals (*F*_*ST*_ values, in panel a), the difference in heterozygote frequencies (panel b), and log transformed P values for associations on LG3 (panel c). Panel (d), for the NMP and flanking regions, shows the associations and the minimum numbers of recombinant haplotypes at different positions represented as both the genetic map locations in centiMorgans, and the physical positions of scaffolds.

### Differentiation, heterozygosity, linkage disequilibrium, and historical recombination in the NMP region

To investigate the levels of differentiation between the NMP and the MP individuals in the NMP region, we performed several correlated analyses (to test for consistency of the results): we estimated *F*_*ST*_, compared levels of heterozygosity between both phenotypes, and performed a linkage disequilibrium analysis (LD). The NMP region showed strong differentiation between NMP and MP individuals for the associated SNPs, reflected in *F*_*ST*_-values for individual markers as high as 0.7 (Fig. 4.a). Equally, there was strong LD between the 36 associated SNPs (Fig. 5), although interspersed with areas of low LD, among non-associated SNPs. In concordance with the association results, we also found higher levels of heterozygosity in NMP individuals than in MP individuals (average heterozygosity in the NMP region of 0.50 vs. 0.33, respectively, *P* < 0.0001; Figure 4.b). This difference in heterozygosity is specific to the NMP region: in the rest of the genome, the NMP and MP individuals did not differ in heterozygosity (the means were 0.34 and 0.32 respectively, *P* = 0.149). However, heterozygosity of four, supposedly inbred MP individuals was only half of the population average (Supplementary Material S3).

**Figure 5.**
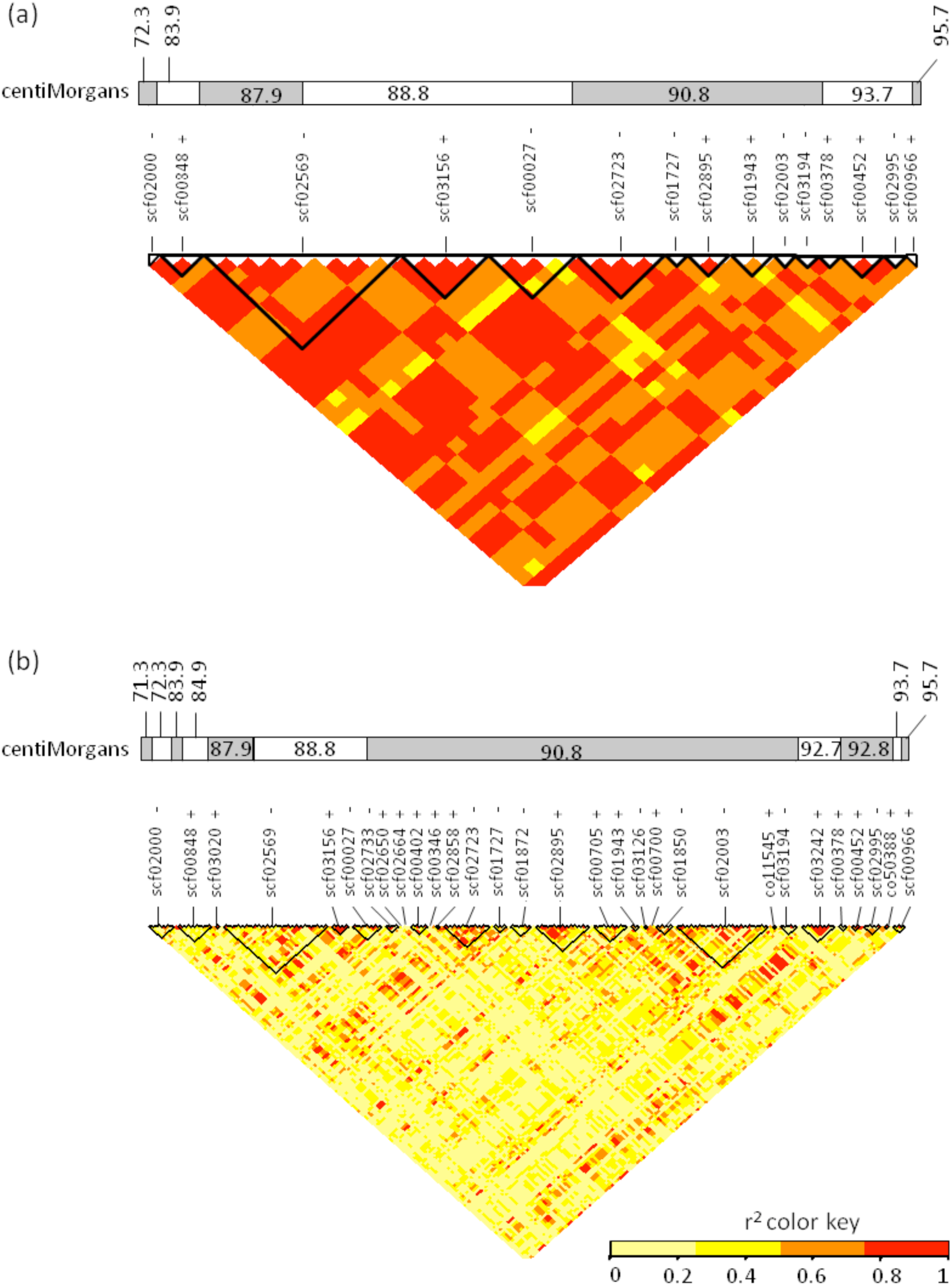
Linkage Disequilibrium Heatmap (r^2^ coefficients) in the NMP region. Results are shown for (a) the NMP associated SNPs, (b) all SNPs mapping to the NMP and flanking region corresponding to the NMP region using the 72 individuals (55 MP, 17 NMP). Black triangles represent scaffolds, and the bi-colored band represents the genetic map positions of each SNP, in centiMorgans. + and -- represent the orientations of the scaffolds (forward and reverse respectively).

Second, to investigate whether the occurrence of multiple, highly associated SNPs distributed across a large region can be explained by a lack of recombination, we tested for historical recombination in the region using Hudson’s four gamete tests (see Material and Methods). These tests detected the presence of recombination in 37 pairs of adjacent polymorphic sites in the NMP region. These historical recombination events were distributed across the entire NMP region, and were found both between and within scaffolds (Fig. 4.d.). This implies that a low level of recombination or gene conversion occurs between the incipient Z and the incipient W chromosome, or has at least occurred historically.

### Identification of candidate genes involved in sex determination, and prediction of the effect of SNP substitutions on amino acid identity

To search the NMP region for genes known to be involved in sex determination / differentiation in other taxa, we analysed the 283 non annotated protein sequences that where located in the NMP region. Of these, 184 returned a BLAST result, but 39 were described as hypothetical proteins in the *D. pulex* draft transcriptome DAPPUDRAFT and did not reliably match any proteins in the NCBI non-redundant protein database. Among the remaining 145 successfully annotated sequences, 121 (82%) had a top hit on *D. pulex* or *D. magna* sequences, while the others matched various arthropods (15 sequences), and other invertebrates (ten sequences). We compared the 145 annotated genes with the NCBI list of 601 genes known to be involved in sex determination or sex differentiation in other invertebrates, and identified 14 candidate genes (Table 3). These include genes involved in sex-specific endocrine signaling pathways, such as a membrane androgen receptors (*zip9*), a tissue specific modulator of the ecdysone response in *Drosophila* (*Broad* complex; Karim et al. 1993), as well as a member of the aldo-keto reductase family (Penning et al. 2000). Scf02569 also harbors a gene resembling transcription factor *sox9*, which acts to inactivate the female differentiation pathway and promote spermatogenesis in males in mammals. In addition, genes involved in chromatin remodeling (lysine-specific histone demethylases, histone deacetylases) are also present in the region. The 14 genes are located on 6 different scaffolds, with scf02569 containing eight of these genes, whereas other scaffolds harbor at most two of these genes (Table 3).

**Table 3.**
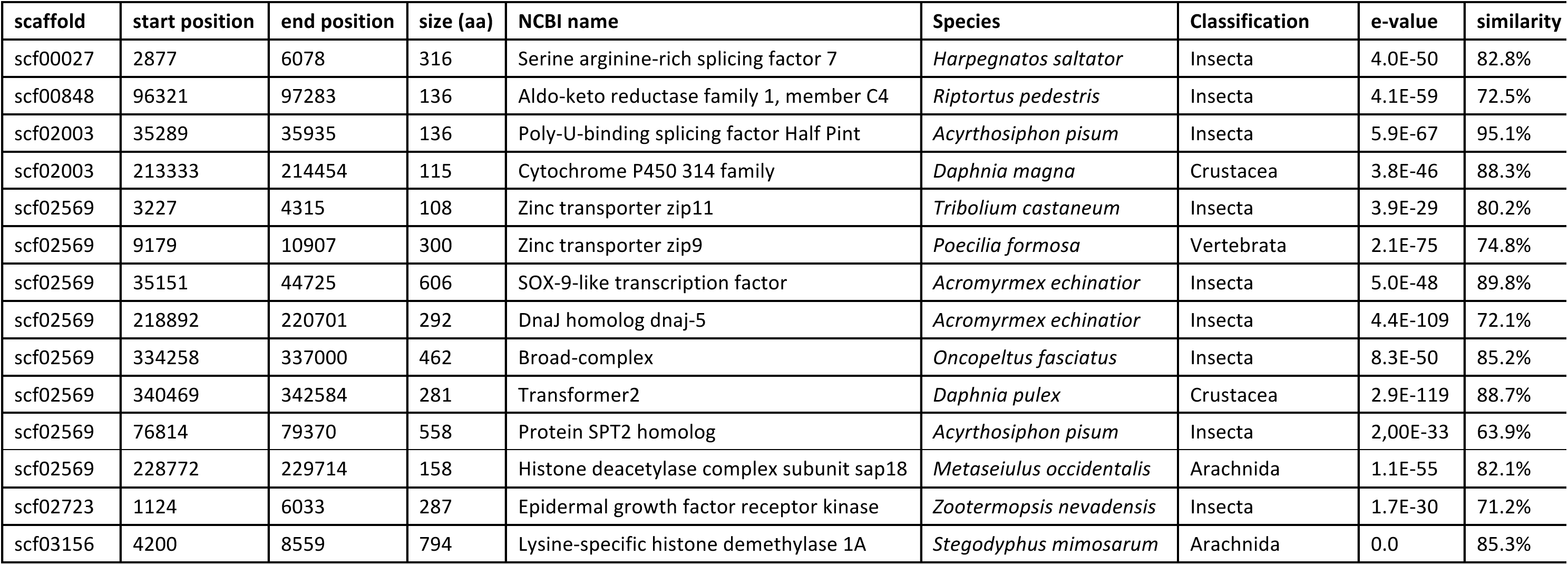
Results of the Blast run showing the 14 candidate genes and their location on the *D. magna* genome. The table reports the start and end position of the gene on the scaffold it maps to, the size of the expected protein (number of amino-acid), the NCBI attributed gene name, the taxon with the best blast hit and its classification, the corresponding e-value, and the percentage of sequence similarity.

| scaffold | start position | end position | size (aa) | NCBI name | Species | Classification | e-value | similarity |
| --- | --- | --- | --- | --- | --- | --- | --- | --- |
| scf00027 | 2877 | 6078 | 316 | Serine arginine-rich splicing factor 7 | *Harpegnatos saltator* | Insecta | 4.0E-50 | 82.8% |
| scf00848 | 96321 | 97283 | 136 | Aldo-keto reductase family 1, member C4 | *Riptortus pedestris* | Insecta | 4.1E-59 | 72.5% |
| scf02003 | 35289 | 35935 | 136 | Poly-U-binding splicing factor Half Pint | *Acyrtosiphon pisum* | Insecta | 5.9E-67 | 95.1% |
| scf02003 | 213333 | 214454 | 115 | Cytochrome P450 314 family | *Daphnia magna* | Crustacea | 3.8E-46 | 88.3% |
| scf02569 | 3227 | 4315 | 108 | Zinc transporter zip11 | *Tribolium castaneum* | Insecta | 3.9E-29 | 80.2% |
| scf02569 | 9179 | 10907 | 300 | Zinc transporter zip9 | *Poecilia formosa* | Vertebrata | 2.1E-75 | 74.8% |
| scf02569 | 35151 | 44725 | 606 | SOX-9-like transcription factor | *Acromyrmex echinator* | Insecta | 5.0E-48 | 89.8% |
| scf02569 | 218892 | 220701 | 292 | DnaJ homolog dnaj-5 | *Acromyrmex echinator* | Insecta | 4.4E-109 | 72.1% |
| scf02569 | 334258 | 337000 | 462 | Broad-complex | *Oncopeltus fasciatus* | Insecta | 8.3E-50 | 85.2% |
| scf02569 | 340469 | 342584 | 281 | Transformer2 | *Daphnia pulex* | Crustacea | 2.9E-119 | 88.7% |
| scf02569 | 76814 | 79370 | 558 | Protein SPT2 homolog | *Acyrtosiphon pisum* | Insecta | 2,00E-33 | 63.9% |
| scf02569 | 228772 | 229714 | 158 | Histone deacetylase complex subunit sap18 | *Metaseiulus occidentalis* | Arachnida | 1.1E-55 | 82.1% |
| scf02723 | 1124 | 6033 | 287 | Epidermal growth factor receptor kinase | *Zootermopsis nevadensis* | Insecta | 1.7E-30 | 71.2% |
| scf03156 | 4200 | 8559 | 794 | Lysine-specific histone demethylase 1A | *Stegodyphus mimosarum* | Arachnida | 0.0 | 85.3% |

Nine of the 176 SNPs identified by RAD-sequencing and mapping in the NMP region were located in genes within which the exon-intron structure could not be determined (very likely because of errors in the assembly or gene structure definition), 69 were in intergenic regions, 17 in 5’ UTR regions, 8 in 3’ UTR regions, 19 in introns, and 53 within a gene (Supplementary Material S2). Of these 53 coding SNPs, 32 of are in non-annotated genes, including one (on scaffold02003 in position 98755) that introduced a stop codon into a gene which was annotated as “hypothetical protein” in the *D. pulex* draft genome. 23 SNPs were synonymous and 30 were non-synonymous mutations. Only one significantly associated SNP induced a non-synonymous mutation in a candidate gene: the SNP at position 4433 on scf03156, inducing a change from Valine to Leucine in the lysine specific histone demethylase. However, RAD-sequencing covers only a small fraction of the genome (here 154745 loci were mapped, with reads of 95bp, representing an estimated 6.12% of the genome). Hence, additional non-synonymous SNPs may be present in parts of the NMP region not covered by RAD-sequencing reads. The same applies for potential regulatory SNPs.

## Discussion

Our results suggest that the NMP phenotype is determined by a single, large genomic region located on LG3. The few significantly linked SNPs on other linkage groups may be in linkage disequilibrium with the major region on LG3, maintained by pleiotropic effects. Alternatively, they may be explained by errors in the genetic map or in the genome assembly, and might, in fact, be variants within the NMP-region. The NMP phenotype mapped to the same LG3 region in all three crosses, involving populations with divergent mitochondrial lineages (Galimov et al. 2011). This indicates either a single evolutionary origin of NMP in *D. magna* or parallel evolution repeatedly involving the same genomic region. Given the divergent mitochondrial sequences and the co-inheritance of mtDNA with the female-determining allele, a single evolutionary origin of NMP in *D. magna* would imply that the female-determining mutation is old. However, parallel (convergent) evolution remains possible, particularly as the NMP region contains more than one gene potentially involved in sex determination and sex differentiation. These genes could represent different mutational targets for NMP-inducing mutations, and hence the mutation may not be the same in all populations, despite occurring in the same genomic region. Finally, rare paternal transmission of mitochondria or transmission of the female-determining mutation through rare males cannot be ruled out, but are unlikely (Galimov et al. 2011; Svendsen et al 2015). Interestingly, a dominant NMP phenotype has also been described in *D. pulex* (Innes and Dunbrack 1993) and in *D. pulicaria* (Tessier and Caceres 2004). Considering that an estimated 150 Myr separate *D. magna* and *D. pulex* (Kotov and Taylor 2011), parallel evolution of the NMP phenotype seems more likely for these two species, rather than a femaleness allele having been maintained for such a long evolutionary time.

According to the classical model of sex chromosome evolution discussed in the introduction, the establishment of a female-determining mutation is followed by additional mutations with sex antagonistic (SA) effects occurring at nearby loci. Recombination between these loci and the female sex-determining locus is deleterious, and hence SA selection is thought to favour a reduction of recombination between the incipient Z and W (or X and Y). Such recombination reduction in the heterogametic sex (compared to recombination in the homogametic one) is considered to be a hallmark of early sex chromosome evolution. Alternatively, the sex-determining mutation may occur in a region with already low recombination. Such regions occur on autosomes, for instance due to inversions or vicinity to the centromere (Hoffman and Rieseberg 2008; Ironside 2010; Joron et al. 2011). If such a region contains, from the outset, multiple linked potential target loci for SA mutations, then a further reduction of recombination after the occurrence of the sex-determining mutation might not be favoured by selection, at least not initially. In our study, the sex-determining mutation mapped to the peri-centromeric region of LG3, which, as expected, has a low recombination rate not only in MPxNMP crosses, but also in MPxMP crosses. Interestingly, the sex-determining mutations of *Papaya* and *Populus* are also found in peri-centromeric locations (Yu et al. 2007, Kersten et al. 2014). In addition, we did not find strong evidence for a further reduction of recombination between the incipient Z and the W chromosome, suggesting that SA selection did not act strongly (if at all) to further extend the low-recombination region containing the female-determining mutation. Yet, SA selection may still play a major role in determining the NMP phenotype. Under the hypothesis that SA selection occurs, multiple loci should contribute to the NMP phenotype, not only for the determination femaleness, but also for the expression or fitness of the female phenotype, or for enhancing maleness of ZZ individuals. Here, we found highly associated SNPs distributed on a total of 3.6 Mb, but separated by regions of low LD in which there is strong evidence for historical recombination or gene conversion. This suggests a role for selection to be acting on several genes in the NMP region. This selection is likely sex-antagonistic, as it appears to maintain different W-linked vs. Z-linked alleles in several sub-regions independently (and thereby maintain the high LD between associated SNPs despite historical recombination). We identified 14 genes in the NMP region that are known to be involved in sex determination/sex differentiation in other taxa. These genes might be potential target for SA mutations. Hence, the NMP region shows all characteristics of a supergene (Joron et al. 2011), with different alleles appearing to be selectively maintained between Z and W in several genes throughout the region. Such a “sex-super gene” is nothing else than the first step towards the evolution of a sex chromosome, even though low recombination in this region is not due to secondary suppression of recombination, but due to a chance event (occurrence of the sex-determining mutation in a large region with pre-existing low levels of recombination). However, such a chance event may, in fact, be one of the possible mechanisms leading to initial suppression of recombination between incipient sex chromosomes.

The transition from ESD to GSD is thought to happen rapidly in nature. However, intermediate stages (such as gynodioecy, or partial GSD) can be evolutionary stable, depending on their origin and genetic control (Charlesworth and Charlesworth 1978). The NMP mutation might first have been favoured by selection because it leads to obligate outcrossing (Innes and Dunbrack 1993). This is supported by the fact that inbreeding depression in *Daphnia* is strong (e.g., Lohr and Haag 2015) and that within-clone mating occurs at an appreciable frequency (e.g., 5 % of MP individuals in our association study were offspring of within-clone mating). Once it attained an intermediate frequency, it was probably also affected by negative frequency-dependent sex-ratio selection, which may have set the stage for additional mutations to gradually improve maleness of ZZ individuals (Innes and Dunbrack 1993; Galimov et al. 2011). The coexistence of ESD and GSD individuals in some populations of *D. magna* might be evolutionary stable. This stability could be due to their reproductive cycle, called cyclical parthenogenesis, which might prevent further evolution towards full GSD. Indeed, only females participate in parthenogenetic reproduction. Hence, in order to profit from parthenogenetic multiplication, all genotypes have to be females at some point during the seasonal cycle. ZZ females might increase their male function during sexual reproduction by prolonging their investment in parthenogenetic clutches (late season parthenogenetic clutches are usually male, Galimov et al. 2011). It is also possible that the maleness of ZZ individuals may have been quantitatively improved by linked genes with SA effects (see above). Nonetheless, at the very end of the season, it may still be more profitable for these females to engage in sex rather than to abandon reproduction or to parthenogenetically produce additional males which might not have the time to develop into adults. If however, *Daphnia* was to complete the transition from ESD to pure GSD (as envisaged by the theory outset in the introduction), the incipient Z homolog of the NMP region would need to acquire a mutation determining the male sex (i.e. a recessive female-sterility mutation), and the two chromosomes would then be called proto-sex chromosomes. This has not happened in *D. magna*, as the ZZ individuals still have ESD and still contribute to the diapause stage production also through female function. However, while hatchlings from diapause stages always seem to be female in *Daphnia*, male hatchlings do occur in the related *Daphniopsis ephemeralis*, living in short-lived environments (Schwartz and Hebert 1987). Hence, it is not inconceivable that some *Daphnia* populations or species might evolve full GSD, perhaps especially under conditions that reduce the importance of the pathenogenetic phase.

The molecular mechanisms underlying male differentiation in *D. magna* and its relationship with ESD are the object of much research (Kato et al. 2008, 2010, 2011; LeBlanc and Medlock 2015). To date, we know that a homolog of the *Doublesex* gene (*dsx*) is found in *D. magna (dapmadsx)* and that it is a major effector regulating the male phenotype (Kato et al. 2011; Salz 2011; Beukeboom and Perrin 2013). The regulation of its expression appears, however, to be different in *Daphnia* compared to other arthropods, as no evidence for sex-specific splicing was identified for the *transformer* gene in male versus female embryos (Kato et al. 2010, 2011; Verlhulst et al. 2010). Instead, the expression of *dapmadsx* might be controlled by an “on/off” mechanism whose activation may involve elements responsive to juvenile hormone (Kato et al. 2011). Our results show that the NMP region does not contain *dapmadsx* nor *transformer*, and hence neither of these genes is the likely location of the female-determining mutation. However, the NMP mutation does contain multiple genes that are potentially members of the same pathways. These genes include *transformer2 (tra2)* on scf02569, a splicing regulator known to interact with *transformer* to control the female sex-specific splicing of *dsx* in insects. It will be interesting to investigate if and how *tra2* interacts with *dapmadsx.* Moreover, we found four genes involved in hormonal pathways (*zip9*, *zip11*, *Broad* complex, aldo-keto reductase), which might be relevant as *dsx* expression might be controlled by hormone-responsive elements. Furthermore, scf02569 contains a gene resembling transcription factor *sox9*, which inactivates the female differentiation pathway and promotes spermatogenesis in males mammals. Overall, the presence of these 14 genes in the NMP region is congruent with a major role of this region in sex determination/differentiation. In addition, scf02569 is a strong candidate for carrying the original sex-determining mutation. Not only does it contain 25% of the significantly associated SNPs and eight of the 14 candidate genes, but one of these candidate genes (*tra2*) appears to be one of the most promising ones, based on its function in other organisms, and on what is currently known about the mechanisms underlying male differentiation in *D. magna*.

In the present study, we determined that the dominant female sex-determining locus observed in multiple population of *D. magna* maps within a single genomic region in the peri-centromeric region of LG3. With a pre-existing low recombination rate, the region contains multiple genomic regions in high LD spanning a total of 3.6Mb, separated by regions of low LD due historical recombination or gene conversion likely occurred. The strong LD is likely maintained by sex antagonistic selection, since multiple genes involved in both males and females sex differentiation were found throughout the region. We conclude that *D. magna*’s LG3 carries a sex-related supergene, and is an incipient W chromosome, the youngest stage that might be possible to empirically observe in sex chromosome evolution. *D. magna* is thus a very promising model for studying the evolutionary transitions from ESD to GSD and early stages of sex chromosome evolution.

## Material and Methods

### Genetic linkage mapping of the NMP-determining region, and assessment of recombination rates

In order to map the genomic region responsible for the NMP phenotype, we performed experimental crosses between known NMP and MP genotypes. Because the NMP locus is believed to be heterozygous dominant in females, the goal was to maximize heterozygosity of the mother NMP clone, while ensuring the MP father was homozygous or had different alleles. All three crosses involved outcrossing between two populations to ensure that a sufficient number of markers had different genotypes between fathers and mothers. One of the crosses also involved a NMP mother that was already a hybrid between two populations, again in order to maximize heterozygosity. The first cross called “NMPxMP_1” involved a NMP female from Volgograd, Russia (48.53 N, 44.486944 E) and a male from Orog-Nur, Mongolia (45.032708 N, 100.718133 E) as well as 66 of their F1 offspring. The second cross (“NMPxMP_2”) used a hybrid NMP female, which was produced by crossing a NMP female from Moscow, Russia, (55.763514 N, 37.581667 E), with a male from Orog-Nur. The cross then involved this hybrid female and a male from Vääränmaanruskia, Finland (60.271617 N, 21.896317 E), as well as 54 of their offspring. The third cross (“NMPxMP_3”) involved a NMP female from Yakutsk, Russia (61.964047 N, 129.630956 E) and a male from Rybnoye, Russia (56.425003 N, 37.602672 E), as well as 22 of their offspring. The three NMP females had divergent mitochondrial haplotypes (based on a 534 bp alignment of the COI sequences, the average number of substitutions per site was 0.037 between Yakutsk and Moscow, 0.041 between Yakutsk and Volgograd, and 0.010 between Moscow and Volgograd; Galimov et al. 2011; Genbank accession number JF750770.1 for Moscow, AY803073.1 for Volvograd, and [added upon acceptance] for Yakutsk). Microsatellite markers distributed across the genome were used to investigate the parental lines. Markers that were heterozygous in the NMP mother and for which father and mother had different genotypes were selected and genotyped in the offspring. For all these markers, it could unambiguously be determined which of two maternal alleles was transmitted to a given offspring. Hence, linkage of markers to the NMP-determining region could be assessed, by assaying co-transmission of maternal alleles with the phenotype (i.e. with the W vs. Z chromosome). Before genotyping, the offspring were thus phenotyped using the juvenile hormone Methyl Farnesoate, which triggers the production of males in MP strains but not in NMP strains of *Daphnia* (Galimov et al. 2011).

Microsatellite loci were amplified using the M13-protocol (Schuelke 2000): For each locus, unlabeled forward and reverse primers were used together with fluorescently labelled, universal M13 primer. The forward primer consisted of a locus-specific part as well as an overhang complementary to M13. PCR reactions were carried out using the Type-it Microsatellite PCR Kit (Qiagen) according to the manufacturer’s protocol with an annealing temperature of 60°C. After 22 cycles, the annealing temperature was lowered to 53°C for another 20 cycles in order to allow for proper M13 annealing. The resulting PCR products were diluted four times and mixed with a LIZ5000 size ladder (Applied Biosystems). Samples were genotyped using ABI 3730 capillary sequencer and GENEMAPPER software v. 3.0 (Applied Biosystems). A total of 81 microsatellite loci (see Supplementary Material S1) were tested in the parents. Of these, 60 were polymorphic in one or both parents and thus genotyped in the offspring (47 in NMPxMP_1 and 21 in NMPxMP_2, partially overlapping). Linkage to the NMP phenotype was assessed with a Fisher’s Exact tests (two-tailed). Some of the markers were specifically designed in regions for which linkage to the NMP phenotype was suspected based on information from an earlier version of the genetic map (Routtu et al. 2010; Routtu et al. 2014) and the initial finding of weak but significant linkage of one marker (dm_scf00243_208642) in the NMPxMP_1 cross. Therefore, the markers do not represent a random sample throughout the genome. The NMPxMP_3 cross, which included the NMP female with the most divergent mitochondrial haplotype, was done at a later stage. It was only used to test whether NMP maps to the same genomic region as in the two other crosses. Hence, only three loci closely linked to NMP in the first two crosses were also genotyped in the offspring of this cross.

Genetic linkage mapping of the NMP region was carried out in R/qtl (Broman et al. 2003). It soon became evident that NMP mapped to a region of LG3 of the *D. magna* reference genetic map (Dukič et al. in press), of which a first version called v4.0.1 was published in Svendsen et al. (2015). Hence, map construction was done using markers that either showed significant linkage with the NMP phenotype (*P* < 0.01 in pairwise Fisher’s exact tests) or were found on scaffolds of the *D. magna* genome v2.4 (bioproject reference PRJNA298946, on the NCBI repository: https://www.ncbi.nlm.nih.gov/bioproject/?term=PRJNA298946) that had been mapped to LG3. Markers that had more than one third of missing genotypes (amplification failures, etc.) were discarded.

The mapping procedure consisted in first positioning and ordering the markers according to previously available data (Dukič et al. in press), if possible, and secondly by estimating genetic distances among the ordered markers according to standard procedures. Specifically, we ordered our markers by using the cM position of the nearest mapped SNP from the same scaffold in the reference genetic map v4.0.1. Microsatellite markers on scaffolds that were not mapped in the reference genetic map were positioned according to the estimated recombination fraction in our crosses between these markers and already mapped markers. The only exception to this procedure was done for microsatellite marker scf02066_483524, which is located on a mis-assembled part of scf02066, closely linked to the end of scf00494 (Dukič et al. in press), and thus was ordered according to this position. Once ordered, Kosambi-corrected genetic map distances among all markers were recalculated from the offspring genotypes of our crosses using R/qtl (with the option sliding-window = 8 markers).

To test for a reduction of recombination around the NMP region in the MPxNMP crosses versus the reference MPxMP cross of the genetic map v.4.0.1, we compared the genetic distances for intervals of adjacent markers between the crosses. Specifically, for each interval, we assessed the number of recombinant vs. non-recombinant individuals in each of the NMPxMP crosses versus the reference cross, and tested for significant differences using Fisher’s Exact tests (two-tailed) implemented in R core package stats (R Development Core Team, 2008).

### Association mapping of SNPs in the NMP region

To further characterize the NMP region, we used single-nucleotide polymorphism (SNP) data obtained by RAD-sequencing of a random sample of 72 individuals (17 NMP and 55 MP) obtained by hatching resting stages from the Moscow population. The details of the RAD-sequencing protocols, alignment to the *D. magna* reference genome, and SNP calling are given in Supplementary Material S4. We obtained 7376 SNPs to be analyzed (Supplementary Material S2 for an excel document containing all SNPs). We first performed a genome-wide association study, using the expectation that any bi-allelic SNP functionally related to NMP or tightly linked to it should be heterozygous in NMP individuals and homozygous for the more frequent of the two alleles in MP individuals, corresponding to the ZW and ZZ genotypes, respectively. Only sites with a minor allele frequency larger than 0.1 and less than one third of the individuals with missing genotypes were used in the analysis. For each retained bi-allelic site, we grouped individuals into four categories: Heterozygous NMP individuals, homozygous NMP individuals, homozygous MP individuals for the major allele, and MP individuals non-homozygous for the major allele (either heterozygous or homozygous for the minor allele). For each site, we counted the number of individuals in each of the four categories and calculated the expected number of individuals (under the null-hypothesis of no association) using standard Hardy-Weinberg proportions with allele frequencies estimated across all individuals. We then used Pearson’s Chi-square tests with two degrees of freedom to evaluate the genotype-phenotype association at each site. However, in order to only test the hypothesis specified above, any excess that went in the direction opposite the hypothesis (for instance an excess of non-heterozygous individuals among NMP) was discarded (i.e., was not taken into account for the overall Chi-square value). The R script of this association analysis is given in Supplementary Material S5. The significance of the association was assessed by correcting the *P*-value of the Chi-square test according to an overall false discovery rate (FDR) of 10^−5^ using the p.adjust function of the R core stats package.

### Differentiation, heterozygosity, linkage disequilibrium, and historical recombination in the NMP region

To confirm that the NMP associated region is highly differentiated between NMP and the MP individuals, we estimated *F*_*ST*_ for each SNP with the R package PopGenome (Pfeiffer et al. 2014). We also performed a linkage disequilibrium (LD) analysis of the NMP region and investigated relative levels of heterozygosity for MP and NMP individuals. Note that all three analyses test essentially for the same thing (Charlesworth et al 1997): If there are many strongly associated SNPs, we would expect *F*_*ST*_ to be high, heterozygosity to be high in NMP individuals, and LD to be high, at least between associated SNPs. All three analyses were run to check for consistency, and in the case of heterozygosity also as a comparison with other genomic regions. The subsequent analyses were restricted to cM positions between 85 and 95 of the genetic map, which includes the NMP region as well as some flanking regions on either side. However, due to the dearth of recombination in this region, the relative position and orientation of many of the scaffolds is unknown (several entire scaffolds having the exact same cM position). Hence, we first inferred the likely relative position and orientation of these scaffolds by LD mapping in MP individuals, assuming no structural rearrangement of those scaffold between MP and NMP individuals (see Supplementary Material S6 for details of the LD mapping procedures). Once the inferred physical map was established, we used it to plot the genotype-phenotype association in the region as well as *F*_*ST*_, and heterozygosity.

Linkage disequilibrium for all individuals (i.e., including NMP) was then estimated via pairwise *r*^*2*^ for each pair of SNPs (see Supplementary Material S7), using the program MCLD (Zaykin et al. 2008). Significance was tested using 9999 permutations, and the extent of LD was visualized using a heatmap constructed in R using the LDheatmap package (Shin et al. 2006). To test for signatures of historical recombination between W and Z haplotypes, we used a filtered dataset composed of 140 polymorphic sites in the region, retaining just one site per read (a maximum of two polymorphic sites on the same read were present in the whole data set, but SNPs on the same read were always in full linkage). We then phased these data using the GERBIL program implemented in the package GEVALT V2.0 (Davidovich et al. 2007). We did not allow the program to infer missing genotypes, since the algorithm favors the use of the more common allele, here the Z allele (only 17 out of 144 haplotypes are W haplotypes). The phasing resulted in two Z haplotypes for each MP individual and in one W and one Z haplotype for each NMP individual (Z haplotypes were identified according to the allelic state of the two most strongly associated SNPs named scf2723_2194 and scf2723_13482). Because we could not exclude genotyping nor phenotyping error (the latter only in the direction of falsely identifying an individual as NMP), we used conservative criteria for the test: we first removed the NMP individual (RM1-01) which had a high genotypic resemblance to MP and thus resulted the highest evidence for recombination (this individual was most likely mis-phenotyped during the hormonal exposure). Furthermore, before carrying out the test we corrected singleton variants within each haplotype group: if a variant was present in only one haplotype in the group, we reverted its state to the majority allele in this group. Overall, 27 loci and 6 loci out of 140 loci were reverted in W and Z, respectively. This conservatively assumes that all these singleton variants were due to genotyping error (note that loci with a minor allele frequency of < 0.1 across both groups had already been excluded during the initial filtering, see Supplementary Material S8 for the list of uncorrected and corrected haplotypes). Finally, to test only for recombination between Z and W haplotypes (as opposed to recombination within Z), we performed the Hudson’s four-gamete test, which stipulates that the presence of the four possible gametes in a pair of segregating bi-allelic sites is a sign of historical recombination or gene conversion. We inspected all instances where recombination was detected by the test and retained only those where the inferred recombination had occurred between the Z and the W haplotypes.

### SNP effect and identification of candidate genes involved in sex determination

To assess whether the NMP region contains candidate genes with known functions related to sex differentiation or sex determination, we extracted all 1306 protein sequences corresponding to transcript sequences located on the scaffolds in the NMP region, and reduced isoform redundancy using BlastClust (http://toolkit.tuebingen.mpg.de/blastclust) with the following parameters: minimum length coverage of 60%, minimum identity of 90%, minimum transcript size of 100 amino acids. This resulted in a set of 361 protein sequences, which we trimmed by hand to remove further the redundancy (we only kept one transcript for each gene). The retained 283 protein sequences (see Supplementary Material S9 for the complete list) were blasted against the NCBI nr database with Blast2GO (Conesa and Götz 2008), using blastp and a maximum e-value of 10^−10^. Annotated genes were then compared with a list of 601 genes obtained from the NCBI gene data base using the keywords “sex determination” and “sex differentiation”. We also used the GFF file (also available on NCBI, bioproject reference PRJNA298946), which contains gene features of *D. magna*, to classify each SNP in the NMP region according to whether it induces a synonymous or a non-synonymous change. This analysis was done using the software tool IGV (Robinson et al. 2011).

## Acknowledgements

We thank the Zoo of Moscow and N. I. Skuratov for sampling permits, and David Frey for help with culture maintenance indoors. We thank the Department of Biosystem Science and Engineering of the ETH Zurich, in particular C. Beisel and I. Nissen for Illumina sequencing, and we gratefully acknowledge support by M-P. Dubois, the platform Service des Marqueurs Génétiques en Ecologie at CEFE, and the genotyping and sequencing facilities of the Institut des Sciences de l’Evolution-Montpellier and the Labex Centre Méditerranéen Environnement Biodiversité (CeMEB). We thank M. Roesti for the modified RAD-seq protocol and for the discussions and scripts on the linkage disequilibrium analysis. We thank the University of Fribourg, the Montpellier Bioinformatic Biodiversity plateform and the Labex CeMEB for access to high-performance computing clusters. The sequence data for the *D. magna* genome project V2.4 were produced by The Center for Genomics and Bioinformatics at Indiana University and distributed via wFleaBase in collaboration with the *Daphnia* Genomics Consortium (project supported in part by NIH award 5R24GM078274-02 “*Daphnia* Functional Genomics Resources”). We also thank Peter Fields and Deborah Charlesworth for their constructive comments at various stages of this paper, as well as David Innes and two other anonymous reviewers who contributed to its great improvement during the reviewing process. This work was supported by the Swiss National Science Foundation (Grant no. 31003A_138203), the Russian Foundation of Basic Research Grant 16-04-01579, the European Union (Marie Curie Career Integration Grant PCIG13-GA-2013-618961, DamaNMP). and the Council on Grants from the President of the Russian Federation in support of leading scientific schools (NSh-9249.2016.5).

## Supplementary material

### S1: Table: microsatellites.xls

Excel file giving the list of the 81 microsatellite markers tested in this study.

### S2: Table: snp_data.xls

Excel file containing the list of SNPs obtained from the RAD-sequencing panel for all individuals. Information listed: Linkage Group; order of SNP on the genetic map; CentiMorgan position on the genetic map; basepair position on the physical map; MegaBase position; scaffold where the SNP maps to; position of the SNP on the scaffold (in bp); major allele; minor allele. Additional information: Chi square value; associated *P*-value; mutation location and type (intergenomic, intron, 5’ / 3’ UTR; amino acid substitution); synonymous or non synonymous mutation; gene impacted.

### S3: Figure.

Caption: Distribution of the percentage of relative heterozygosity in MP (black) and NMP (grey) individuals. Calculations were performed without LG3, as this chromosome shows a higher heterozygosity in NMP individuals.

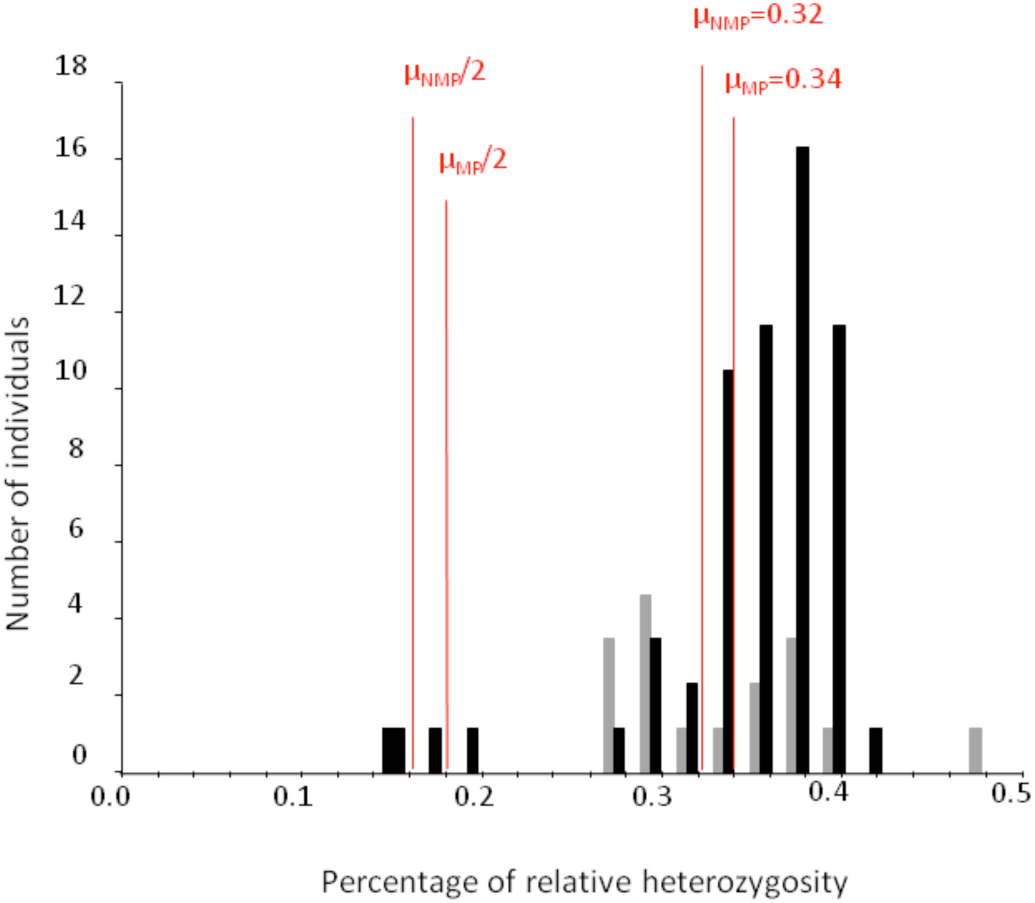

### S4: Text : RAD-sequencing and SNP calling protocol

We used the RAD-sequencing protocol developed by Etter et al. (2011) with a few modifications. The 72 individuals were divided in 2 libraries. Prior to DNA extraction, individuals were treated for 72 hours with three antibiotics (Streptomycin, Tetracyclin, Ampicilin) at a concentration of 50 mg/L of each antibiotic and fed with microscopic glass beads (Sephadex “Small” by Sigma Aldrich: 50 µm diameter) at a concentration to 0.5g/100 mL. The aim of this treatment was to minimize contaminant DNA (i.e., bacterial DNA or algal DNA) in in the gut and on the surface of the carapace. Genomic DNA was extracted using the Qiagen Blood and Tissue kit following manufacturer’s instructions and digested with PstI (New England Biolabs). Digested DNA was barcoded with individual-specific P1 adapters and pooled to create a library containing 2100ng DNA. The pooled library was sheared on a Bioruptor using 2 times 3 cycles (1 cycle 30 seconds ON, 1 minute OFF), and fragments between 300 and 500bp were selected through agarose gel electrophoresis. DNA fragments were blunted and a P2 adapter was ligated. The library was amplified through PCR (30 seconds at 98°C, followed by 18 cycles of 10 sec. at 98°C, 30 sec. at 65°C and 30 sec. at 72°C; a final elongation step was performed at 72°C for 5 min.). A final electrophoresis was performed to select and purify fragments between 350 and 600bp. Each library was sequenced on a single lane of an Illumina HiSeq 2000, using single-end 100 cycle sequencing by the Quantitative Genomics Facility service of the Department of Biosystem Science and Engineering (D-BSSE, ETH), Basel, Switzerland.

The quality of the raw sequencing reads (library-wide and per-base) was assessed with FastQC (http://www.bioinformatics.babraham.ac.uk/projects/fastqc/), and reads were checked for barcode integrity, absence of adapter sequences within the reads, and integrity of the PstI cut site. The reads were sorted individually by barcode and filtered to remove reads with uncalled bases and an overall base quality score of less than 24. Reads were subsequently aligned to the *Daphnia magna* genome (V2.4; *Daphnia* Genomic Consortium, bioproject reference PRJNA298946, on the NCBI repository: https://www.ncbi.nlm.nih.gov/bioproject/?term=PRJNA298946) using BWA v.0.7.10 (Li and Durbin 2009). Reads that did not map to the reference genome or that mapped to more than one place were discarded. The successfully mapped reads were filtered according to mapping quality (end-to-end mapping with a mapping quality score of at least 25, no more than eight high quality substitutions).

Assignment of reads to RAD loci (defined by unique 95 bp locations on the reference genome) and genotype calling was performed in Stacks V1.19 with a bounded SNP model in pstacks (--bound_high of 0.04, according to the base call error rate provided by the sequencing facility) and allowing a maximum of two high frequency haplotypes (i.e. alleles) per locus per individual. Loci with more than two high frequency alleles were excluded because of a too high risk of falsely mapping paralogous reads to a single locus. Cstacks and sstacks were operated with default settings and with the -g option to use genomic location as method to group reads. The distribution of the minor allele frequency indicated that heterozygous loci usually had a minor allele frequency ranging between 0.2 and 0.5 within an individual. We thus fixed the max\_het\_seq parameter to 0.2 in the program genotypes. As such, potentially heterozygous genotypes with a minor allele frequency of between 0.05 (default homozygote cut-off) and 0.2 were considered ambiguous and were scored as missing in the results. Loci were also filtered according to sequencing depth: Loci with less than 20 reads were discarded (to reduce uncertainty in genotype calls) as were reads with a more than five times higher depth than the average depth across individuals (to reduce the risk of including repetitive elements).

After final genotype calling, loci were mapped to the Daphnia magna genetic map v.3.0 (Dukič et al, submitted). This was done by extracting for each RAD locus the linkage group and cM position of the nearest map-markers on the same scaffold and, if needed, by extrapolating the cM position of the RAD locus by linear extrapolation between the two nearest map-markers.

### S5: Script: association.R

Annotated R script providing the function we used to perform the association analysis at the genome wide level. This script comes partly from the one used in Joron et al. 2011, referenced in the article.

### S6: Text. Protocol for the physical ordering of scaffold in the NMP associated non-recombining region of LG3.

The region controlling the NMP phenotype maps to a region with a low recombination rate in the reference genetic map. This results in the fact that numerous scaffolds are mapped to the exact same cM position, and their relative position and orientation amongst each other are not resolved. Hence, no physical order of the scaffolds in the region can be obtained from the genetic map. This is problematic for genome-wide association studies and fine mapping of the NMP locus, especially for determining whether the NMP phenotype maps to one or multiple specific sub-regions. In an attempt to physically order and orientate the scaffold in the NMP region, we performed linkage disequilibrium (LD) mapping, which uses data on LD from a single population and therefore can make use of historical recombination events present in the data. LD mapping relies on the expectations that two physically close loci should show a high correlation in their segregation patterns in a population (and thus high LD), since recombination events between the two loci should be rare. It is a population based method, so that individual discrepancies with the global population pattern cannot be tested.

Because the NMP region on the incipient W chromosome might carry phenotype specific rearrangements, we based the physical ordering using LD mapping only on the 54 MP individuals sampled from the MOS population. We performed LD mapping on a region of LG3 between 85cM and 95cM, in order to use SNPs just outside the NMP linked region as anchor. The MOS dataset contains SNPs on 30 mapped scaffolds in this region. Among these, there are three groups of scaffolds for which the relative position and orientation could not be resolved with the genetic map: two scaffolds at position 88.8 cM, 17 scaffolds at 90.8 cM and 4 scaffolds at 93.7 cM. For physical ordering of the scaffolds within each of these groups, we first calculated *r*^*2*^ values for each pair of SNPs with MCLD (Zaykin 2008), which is based on the correlation of segregation of genotypes in natural populations, avoiding the need to phase the data, but removing any individual particularities in segregation pattern. We then averaged the *r*^*2*^ values of the three terminal SNPs on each side of each scaffold and created a matrix of pairwise average *r*^*2*^ values between each pair of scaffold extremities for each of the cM groups separately. When more than two scaffolds had to be ordered in a group, we perform a hierarchical clustering analysis to identify “starting clusters” (highly linked scaffold extremities), using the hclust function of the R core package stats. Scaffolds were then added one by one to the starting clusters following the hierarchical order obtained from the hclust dendogram, and oriented in a way that maximized the average *r*^*2*^ values between adjacent scaffold extremities.

### S7: Table: LD_nmp_region.xls

Excel document containing calculated *r*^*2*^ values for the SNPs present in the NMP non recombining region, in order to order and orientate scaffolds in the region not recombining in the reference MPxMP cross.

### S8: Table: phased_haplotypes.xls

Excel document containing the list of the raw and the corrected haplotypes phased in this study, along with the SNP coordinates for each position.

### S9: Text: nmp_region_gene_content.fasta

FASTA formatted document listing the 283 genes used in the analysis.

#### Raw genomic data

The FASTQ files of all the individuals used in this study will be available on the SRA database upon acceptance of the manuscript.

